# Monitoring of the invasive round goby in an estuarine seascape based on eDNA

**DOI:** 10.1101/2024.05.06.592655

**Authors:** Leon Green, Thomas G. Dahlgren, Alizz Axberg, Marina Panova, Matthias Obst, Per Sundberg

## Abstract

Non-indigenous species (NIS) is one of the five most global concerns when it comes to ecosystem services and threats to native biodiversity. This is especially true in aquatic environments which are harder to monitor than terrestrial environments, and NIS are often found first when they are fully established and basically impossible to eradicate. In marine environments, this is further complicated due to the connectivity and difficulty of eradication. The development and implementation of effective monitoring methods for marine NIS are therefore crucial to enable early detection useful for management strategies. In this study, we develop and evaluate environmental DNA monitoring using quantitative (q)PCR as a means to assess presence of the euryhaline round goby fish (*Neogobius melanostomus*) in a seascape environment close to Scandinavia’s largest shipping port. We developed a dPCR assay for the species, targeting a region of 12S gene, and verified its specificity compared to other common species from the gobiid-family in the region. We also experimentally determined the decaying rate of round goby DNA in water to a half-life of 9.8 hours in 15 PSU and 15°C with live fish in captivity. Finally, we sampled 10 sites within a 400 km^2^ area for eDNA and presence of the species using fyke-nets and baited remote video to validate the accuracy of the water samples to predict presence, and abundance. We found that the number of DNA copies extracted from the water samples varied strongly at sites where round gobies were caught in nets or on video, but that the average value from four water samples significantly correlated with an average value from four video samples, and also with the total catch at each site. The eDNA assay also detected signals from the species at sites where no fish were caught by fishing or on video. These results show that this method is highly sensitive for the species even in low abundance, and with sufficient amounts of replicates, it can be possible to determine the relative abundance between sites.

## 1.0 Introduction

Non-indigenous invasive species (NIS) has long been acknowledged as a global problem affecting local ecosystems, their species, functions and the people depending on their services (Elton, 1958, Pyšek et al., 2020). In the marine environment, these issues have also long been apparent (Thresher & Kuris, 2004). Unfortunately, the number of species translocated by for example shipping traffic is continuing to expand, projecting the estimates of the number of yearly species invasions to increase by orders of magnitude between 2020 and 2050 (Sardain, Sardain, & Leung, 2019) despite the implementation of the IMO ballast water convention in 2017. The WWF is already including NIS in the top five global threats contributing to biodiversity loss (WWF, 2020), but the severity of ecological and economical impacts of NIS is undoubtedly expected to grow in the future (Anton *et al*., 2019; Cuthbert *et al*., 2021; Haubrock et al 2022). For example, climate change impacts are likely to exacerbate range expansions of invaders (Rahel & Olden, 2018; Giménez et al., 2020) and open up new routes for introductions (Miller & Ruiz, 2014), calling for increased and more holistic management efforts (Bonebreak et al., 2018). This is especially problematic for aquatic environments, since they are difficult to monitor, and efforts of eradication are more costly and time consuming than on land (Thresher & Kuris, 2004). Eradication and control is a lot more efficient and cost-effective during the early stages of a biological invasion characterised by small population sizes and low density (Ahmed et al., 2022; Cuthbert et al., 2022). However, low density also limits detection and monitoring of the species (Bax et al., 2008). Developing tools to increase the detection opportunities and monitoring of NIS is therefore of high interest to stakeholders and conservation actors (Robertson et al., 2020).

Monitoring of species spatial and temporal distributions is important to understand resilience to habitat change or disturbance or, in the case of invasive species, the severity of the invasion and any impact on local ecosystems (Simberloff et al., 2013). Furthermore, it is important to have methods that can detect invasive species in the early phases of introductions if eradication at all will be possible (Early et al., 2016). Traditional sampling techniques for fish include various active and passive gear and more recently also capturing of images and analyses of environmental DNA. Monitoring in complex, fragmented and dynamic habitats, such as archipelagos or estuaries, requires a higher sampling effort to achieve a relevant statistical power, often associated with significant costs (Stockwell & Peterson, 2002; Nygård et al., 2016). This is further complicated by the problem of detecting mobile species at low densities in a large system. Using traditional methods, is very unlikely that a species will be detected before it reaches sufficient numbers to be considered established. At this population density, it is most likely difficult to eradicate or even control the population growth (Roy et al., 2023). Environmental DNA (eDNA) has established itself as a valuable aquatic monitoring tool due to its cost-efficiency (Holman et al., 2019), ability to detect species or their various life history stages that might be hard to visually find (Beng & Corlett, 2020), and from small or emerging population sizes (Doyle & Uthicke, 2021), also for mobile species such as fish (Thomsen et al., 2012). While traditional monitoring methods in several studies outperform eDNA, depending on the question at hand, (Rose et al., 2019; Ulibarri et al., 2017), the method is developing quickly, with promises for uses in invasive species monitoring programs (eg Sepulveda et al., 2020; Wu et al., 2022, but see Danziger & Friederich, 2022).

Challenges remain in understanding how much DNA is released to the environment from an organism (shedding rate), if it varies between species and higher taxa, and how fast eDNA degrades in the environment (degradation rate) and under what biological and abiotic conditions. This is especially important for the above mentioned cases of monitoring invasive species, and underscores the importance of verifying detection efficiency by using different methods (Uthicke et al., 2018).

Studies that include estimates of shedding rates indicate large differences between different types of organisms. Recent research on starfish indicates release rates around 2 million copies of target DNA per minute and individual (Uthicke et al., 2018), cold water corals 114000 copies per minute from a 2 kg colony (Kutti et al., 2020), and fish at around 250000 copies per minute and fish (Maruyama et al., 2014). These studies are all based on captive organisms in laboratory experiments and release rates in the wild may be considerably different. The rate at which eDNA degrades in seawater has also been studied with half-lifes estimated in lab experiments in the range of a day or two (Harrison et al., 2019). However, the concentration of eDNA in marine ecosystems varies considerably in space and time and care must be taken in interpreting results (Rourke et al., 2022).

The value of eDNA in informing invasive species management eventually comes down to the ability to accurately confirm the presence of a target species and to avoid the risk of false negatives and false positives (Rourke et al., 2022). Development and testing of species specific assays for quantitative PCR such as the dPCR (digital PCR) technology is a straight-forward process and for many invasive species assays have already been published and used for monitoring marine indigenous species in Scandinavian waters (e.g. Knudsen et al., 2022). *Neogobius melanostomus* has had a relatively long history of spreading in the Baltic region. To detect the species, a conventional PCR assay targeting the COI gene has previously been developed for use in the central European river systems where the species is also well established (Adrian-Kalchhauser & Burkhardt-Holm, 2016). However, in a seascape, coastal conditions risk deteriorating the DNA fast (Collins et al., 2018). Digital (dPCR) and digital droplet (ddPCR) PCR potentially allow for quantification of very low concentrations of target DNA (Doi et al. 2015; Hunter et al. 2017). Digital PCRs methods such as ddPCR and dPCR provide absolute quantification, i.e. the number of target DNA molecules per μl water, potentially enabling a comparison of eDNA density with other methods of monitoring (Morin et al. 2023).

The round goby (*Neogobius melanostomus*, Pallas 1814) is Scandinavia’s first known invasive marine fish and appeared in the Baltic in the early 1990’s (Kornis et al., 2012). The species originates from the Black Sea region, and populations in Europe have been found to be of different ‘ecotypes’ adapted to either freshwater rivers or brackish/salt water (Green et al., 2021). In 2010, the brackish ecotype of the round goby was found in Gothenburg (Green et al., 2022). Since then, it has also spread into the adjacent archipelago and can now be found across the entire salinity gradient of the river estuary, and seems able to tolerate fully marine and freshwater conditions (Green et al., 2022).

This bottom dwelling, up to >30 cm long fish can have big impacts on the benthic community (Van Deurs et al., 2021) and is important to monitor for efficient targeted reduction efforts (Bradley et al., 2019, Green & Grosholz, 2021; ICES, 2022; McConnachie et al., 2015).

## 2.0 Aims

The study aimed to test the possibility of using eDNA to monitor round goby, and to verify the approach by accompanying other techniques (fishing and video surveys). To understand the probability of false positive results (i.e. temporary eDNA from ballast water discharge) the decaying rate of DNA in the water after the source (the fish) was removed was also measured.

## 3.0 Methods

### 3.1 dPCR assay development

The assay was designed to maximise the sensitivity by targeting mtDNA genes that are present in multiple copies per cell, specifically the 12S gene. As a part of this project, we also tested an assay targeting the COI gene. It showed however a weak positive signal to the painted goby *Pomatoschistus pictus* DNA at the annealing temperature below 69°, and was subsequently abandoned for this reason. For the 12S gene, sequences for *N. melanostomus* as well as the five most abundant sympatric goby species (all native) within the study area (Table 1) were sourced from the NCBI database, and aligned in the Geneious software to detect the region with the highest divergence between the species. For *Pomatoschisthus flavencens* and *P. pictus*, the available partial 12S sequences had very short overlap with the 12S fragments available for other species, and these therefore could not be included in the primer design. The divergence between *N. melanostomus* and the other gobiid species (*G. niger, P. microps* and *P. minutus*) in the 12S region chosen for the primer design was 18-22%. (The divergence level between *N. melanostomus* and *P. pictus* and *P. flavescens* in their overlapping fragments was 15% - 18%). The forward primer, the reverse primer and the probe sequences were designed using Primer3 following Bio-Rad guidelines for the primer/probe assay design (BioRad Bulletin 6407, n.d.). The designed primers and probe (Table 1) had a perfect match to the DNA of *N. melanostomus* and several mismatches to the other gobiid species (Table 2). Potential primer matches to other organisms was tested in the Primer-BLAST tool at NCBI. No perfect matches were found for either forward or reverse primers to any other organism. However, the Primer Blast detected several cases when the primers matched to other organisms with 2-5 mismatches per primer, which could potentially lead to non-specific PCR amplification: gobiids *Gobius paganellus, Callogobius sclateri, Callogobius maculipinnis* and *Vanderhorstia* sp., and several scorpion fishes from the family Cottidae. To ensure that these species would not risk contaminating samples from the wild, the geographic distributions of all these species was sourced in the database FishBase (https://www.fishbase.se) and none were found to be present in the Swedish waters or known as non-native species in Northern European marine environments.

**Table 1.**
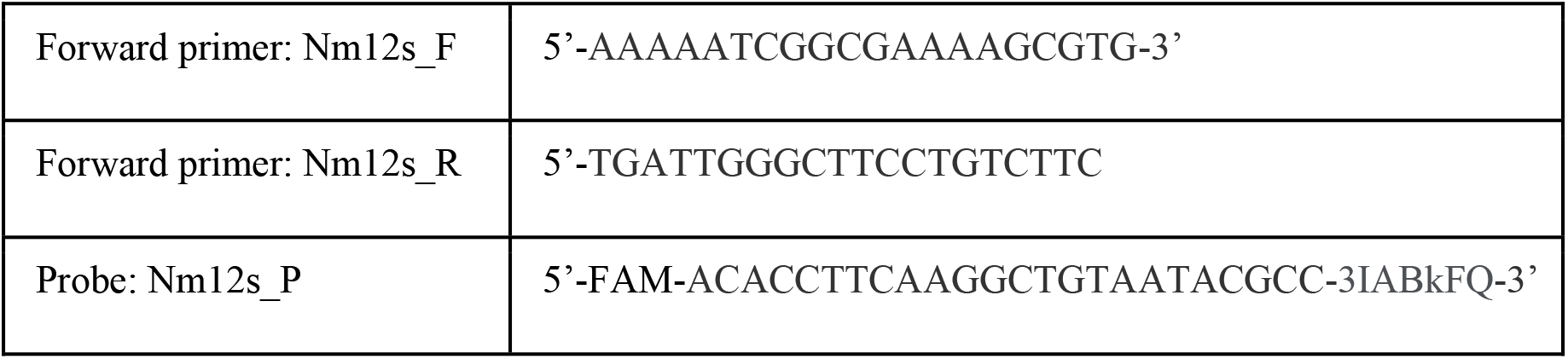
Primer and probe sequences.

**Table 2.** 12S gene sequences used in the primer design and complementary match (number of mismatching positions) between the designed primers and the species DNA sequences for the invasive round goby and the five most abundant sympatric goby species in the sampling area.

| Species | Common name | Accession numbers | Forward primer | Reverse primer | Probe |
| --- | --- | --- | --- | --- | --- |
| *Neogobius melanostomus* | Round goby | 12S:KU755530 | 0 | 0 | 0 |
| *Gobius niger* | Black goby | 12S:EF218630 | 4 | 8 | 5 |
| *Pomatoschistus microps* | Common goby | 12S:MN122842 | 8 | 6 | 5 |
| *P. minutus* | Sand goby | 12S:EF218633 | 8 | 4 | 5 |
| *P. pictus* | Painted goby | 12S: N/A | N/A | N/A | N/A |
| *P. flavescens* | Two-spotted goby | 12S: N/A | N/A | N/A | N/A |

The tests of the designed primers were done using conventional PCR and with DNA samples of *N. melanostomus* and the three common sympatric native gobiid species to assess primer efficiency and specificity (importantly since the primers may amplify the DNA of other species despite several mismatches). In short, tissue samples were collected from euthanized specimens of *N. melanostomus* (N = 1, three replicate DNA extractions), *G. niger* (N=2), *P. minutus* (N=2) and *P. pictus* (N=2) and preserved in 95% ETOH for few days. From these, DNA was extracted using the DNeasy Blood & Tissue kit (QIAGEN), following the manufacturer’s protocol. For the round goby, three replicate DNA extractions were made for a total of 9 DNA samples. Final DNA concentrations of these were measured using a Qubit Fluorometer (Invitrogen) and found to be 1-7 ng/μl. We were not able to obtain samples for *P. microps* and *P. flavescens* for this particular study.

PCR was performed for the *N. melanostomus, G. niger, P. minutus, P. pictus* DNA samples, plus two negative controls: an artificial fish community sample (10 species; no gobiid) and PCR-grade water. The PCR reaction was performed in 25 μL volume, consisting of 12.5 μL KaPa HiFi PCR mix 2x (Roche), 1.25 μL of 12S dPCR primer-probe assay 20x (Bio-Rad), 2 μL DNA template and 9.25 μL nuclease-free water. PCR cycling was conducted at 95 °C for 5 min; 30 cycles at 95 °C for 20 s, 60 °C for 15 s, 72 °C for 15 s min; and 72 °C for 5 min. The dPCR primer-probe assay was ordered from Bio-Rad with primer:probe concentration 900 nm:250 nm and labelled with FAM dye at 5’ and Black Hole Quencher at 3’. In this PCR it was used instead of conventional PCR primers.

As a control for DNA quality, an approximately 260 bp long fragment of the 12S gene was amplified with the universal MiFish primers (Miya et al., 2015). This PCR reaction was performed in 25 μL volume, consisting of 12.5 μL KaPa HiFi PCR mix 2x (Roche), 1 μL each of the forward and reverse primer at 10 μM concentration, 2 μL DNA template and 8.5 μL nuclease-free water. PCR cycling was conducted at 98 °C for 2 min; 30 cycles at 98 °C for 40 s, 62 °C for 30s, 72 °C for 30 s min; and 72 °C for 5 min. The PCR results were visualized on gel electrophoresis (1.5% agarose in 1 x TAE and 5 μl GelRed, run for 30 min at 100V). In the results, all gobiid samples and the fish community sample showed a positive amplification with the universal MiFish 12S-primers (Supplementary material Figure S1a). The 12S *N. melanostomus* assay primers showed strong amplification of the *N. melanostomus* DNA samples but also weak amplification of the *G. niger* DNA and some large non-specific bands (Supplementary material Figure S1b).

To eliminate non-target amplification of the *G. niger* DNA we performed a gradient PCR of the *N. melanostomus* and *G. niger* DNA with the following annealing temperatures: 60°, 62°, 64°, 66°, 68° and 70°C. At annealing temperature ≥64° the PCR band in the *G. niger* sample disappeared (as well as the large non-specific band), while strong amplification of the round goby DNA occurred at all temperatures (Supplementary material Figure S2). From this we concluded that setting the annealing temperature ≥64° ensured only species-specific amplification for this assay.

Finally, the 12S assay was tested on dPCR using QIAcuity One Digital instrument (QIAGEN). Three samples were tested: *N. melanostomus* DNA 0.04 ng/μl; a water eDNA sample from experimental aquaria with the round gobies (see section 3.2 below for experimental details) and a negative control (water). For these three samples, two dPCR reactions per sample were set up in a 26K dPCR nanoplate in 40 μl reaction volumes, containing 10 μl QIAcuity probe Mastermix 4x (QIAGEN), 2 μl primer-probe assay 20x (Bio-Rad), 2 μl DNA sample and 26 μl nuclease-free water. The cycling parameters were 95°C for 2 min, followed by 40 cycles of 95°C for 30 sec and 68.5°C for 1 min (annealing + elongation).

In the results, the 12S assay gave a strong positive signal for both positive samples: approx. 10,000 positive partitions in the *N. melanostomus* DNA sample and approx. 8,000 in the water eDNA samples (out of approx. 25,000 partitions per sample in total), with < 1% variation between the duplicate reactions. No amplification was observed in the negative control. Initially, we chose the high annealing temperature (68°C) to ensure the assay specificity. However, it is above the recommended annealing temperature for dPCR and thus potentially may affect the sensitivity of the assay, especially for environmental samples with very low concentrations of the target DNA. Consequently, additional dPCR tests were performed with the DNA samples from *N. melanostomus* (0.04 ng/μl) and *G. niger* (0.05 ng/μl) and lowering annealing temperatures. In the result we found that even at 64° the assay retains specificity: 1,500 positive partitions (out of approx. 25,000 partitions per sample in total) in the *N. melanostomus* goby DNA sample, none in the *G. niger* DNA and in the negative controls. Therefore, the annealing at 64°C can be applied, especially for environmental samples.

### 3.2 eDNA decay time

To avoid false positives, it is important to know for how long time detectable amounts of DNA remain in the environment after the source (fish) has disappeared from the area (Deiner & Altermatt, 2014). If the DNA has a long “lifespan” it becomes difficult to know if there are living animals of the target species in, for example, a bay, or if there are remnants of old DNA. A tank experiment with captive round gobies was carried out to estimate how long target DNA can be detected in the water after the source (the fish) has disappeared.

Permits for the animal experiments were granted by the Ethical Committee for Animal Research in Gothenburg (permit numbers 5.8.18-03920/2018). Fish were collected from two sites in the harbor of Gothenburg with a mix of methods (overnight cages or baited hook and line). From: the harbor of Arendal (11.8163, 57.6943, N = 14); Klippans färjeläge (11.9103, 57.6920, N = 22). 10 of the fish caught at Arendal were initially intended for another study, and had been acclimated to 20 PSU. An additional acclimation period of 24h during which these fish were ramped down to 15 PSU was therefore done before the animals were introduced to the tanks where the sampling was to be done.

Round gobies were kept in bare tanks (30 liters, untreated tap water mixed with artificial sea salt for 15 PSU ‘natural’ salinity, at 15°C and bubbled with air) for 24 hours before sampling. Three experimental runs with different densities of round gobies (1, 3 and 5 individuals) were conducted over 4 replicates per run. During these runs, the fish were not disturbed and simply left to pass time in the tanks. After 24 hours, the fish were removed from the tanks, and 1 liter of water sampled at ten subsequent times at intervals of six or twelve hours according to the following scheme: T0: 0h; T1: 6h; T2: 12h; T3: 18h; T4: 24h; T5: 30h; T6: 42h; T7: 54h; T8: 78h; T9: 102h; T10; 112 h for a total of 120 samples. Directly after being taken, each sample was filtered through a sterilized 0.22 μm Sterivex filter (Merck Millipore) using one-use 60 ml syringes. After this, each filter was fixed with 95% ethanol inside a labelled 50 ml Falcon tube, and stored frozen at -20°C until processing. After removal from the trial, each fish was euthanized using a lethal dose of MS-222 followed by destruction of the brain and decapitation. They were then weighed to the closest 0.1 gram using a digital scale. From one individual, a 2x2 cm fin clip tissue was also preserved in 95% ethanol as a positive control.

For eDNA sample processing, DNA was extracted from the filters using Nucleospin eDNA water kit (Macherey-Nagel) following the manufacturer’s protocol. The concentration of DNA in the extracts was measured using a Qubit Fluorometer (Invitrogen) The control DNA from the fin clip tissue was extracted with a DNeasy Blood and Tissue Kit (Qiagen) following the methods outlined in Green et al., 2021.

To detect the presence of DNA from the target species in the experimental water samples, the dPCR assay developed in this study was used. dPCR reactions were set up in a 26K dPCR nanoplate in 40 μl reaction volumes, containing 10 μl QIAcuity probe Mastermix 4x (QIAGEN), 2 μl primer-probe assay 20x (Bio-Rad), 10 μl DNA sample and 18 μl nuclease-free water. The cycling parameters were 95°C for 2 min, followed by 40 cycles of 95°C for 30 sec and 64°C for 1 min. Two dPCR reactions were performed for each sample and the average value was calculated. DNA extracted from round goby tissue was used as a positive control in the assay, and DNA/RNA free water as a negative control. As the amount of DNA shed correlates with biomass (Rourke et al., 2021), the results of each replicate were pooled and expressed as copy number per replicate fish mass.

All mathematical calculations were made using R (version 4.1.0, R Core Team, 2021). The equation used to calculate decay rate was:

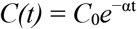

where *C(t)* is the concentration after t hours, *C*_0_ is the concentration at *t*=0, and α is the decay factor.

### 3.3 Sampling sites and metadata for fishing, eDNA sampling

To verify the possibility of eDNA for monitoring of round goby in a seascape, ten sampling sites were selected in the vicinity of Gothenburg for eDNA water sampling, baited underwater video (BRUV) and fishing, during the 8^th^-26^th^ of September of 2021. To avoid cross-contamination between sites through fishing/underwater gear, water samples were taken one week before the fishing started. In addition to the ten near shore sampling sites, eDNA was sampled more offshore from a boat as reference sites and included as a test for how long DNA can travel from expected localities with round goby.

### 3.4 eDNA sampling

At each site (Table 3) four independent samples of water were collected and the eDNA filtered and stored for later analysis. All collection was done using disposable equipment (gloves and sample containers) or in the case of costly sampling equipment - sterilized with 10% bleach solution between each sample. In short, several sub-samples of water were taken using a pole attached 1-litre disposable plastic bucket from the surface, or by using an 1.7-litre Ruttner water sampler (Supplementary figure S2). When sampling with the 1-litre disposable bucket, the sub-samples were pooled into a sterilized two-liter vessel. When depth allowed, the water samples were taken with a Ruttner water sampler near the bottom (except in the case of the reference points, see Table 3 Ref 1 and Ref 2, where the sample was taken in the middle of the water column). Since DNA degradation begins soon after it leaves the source, the 2-litre samples were filtered and fixed within 4 hours of collection following the above described procedure: each sample was filtered through a sterilized 0.45 or 0.22 Sterivex μm filter (Millipore) using one-use 60 ml syringes. Depending on the amount of dissolved matter in the water, the amount of filtered water varied between samples (see Table 3). After this, each filter was fixed with 95% ethanol inside a labelled 50 ml Falcon tube, and stored at -20°C until extraction of DNA. At one of the test sites, 1 litre of DNA/RNA free water was filtered as a negative control.

**Table 3.** Sampling sites for eDNA, BRUV recordings and fishing effort. All sites, number of eDNA samples N = 4, except for reference points where N = 3.

| Site nr | Site name | Position<br>lat N/long<br>E | Salinity<br>‰ | Temp<br>°C | Liters<br>filtered<br>for DNA<br>per<br>sample | BRUV<br>recordings | Fishing Effort |
| --- | --- | --- | --- | --- | --- | --- | --- |
| 1 | Husvik Brännö | 57.634617/<br>11.775452 | 13 | 19.1 | 1 | N = 4 | 3 x 10 meter<br>fyke nets + 6<br>cages |
| 2 | Ferry terminal<br>Vrångö | 57.570120/<br>11.791370 | 16 | 18.0 | 1 |  | 3 x 10 meter<br>fyke nets + 6<br>cages |
| 3 | Stora Amundön | 57.634617/<br>11.775452 | 11 | 18.5 | 1 | N = 4 | 3 x 10 meter<br>fyke nets + 6<br>cages |
| 4 | Ganlets badplats | 57.630130/<br>11.878190 | 7 | 19.7 | 1 | N = 4 | 3 x 10 meter<br>fyke nets + 6<br>cages |
| 5 | Långedrag | 57.670277/<br>11.849265 | 5 | 14.8 | 0.63-1 | N = 4 | 3 x 10 meter<br>fyke nets + 6<br>cages |
| 6 | Hällsvik | 57.699676/<br>11.734819 | 10 | 15.0 | 1 | N = 4 | 3 x 10 meter<br>fyke nets + 6<br>cages |
| 7 | Store Udd<br>badplats | 57.757584/<br>11.794781 | 7 | 17.9 | 0.9-1 | N = 4 | 3 x 10 meter<br>fyke nets + 6<br>cages |
| 8 | Sillviksbadet | 57.741866/<br>11.732092 | 16 | 18.0 | 1 | N = 4 | 3 x 10 meter<br>fyke nets + 6<br>cages |
| 9 | Hönö Klåva | 57.681038/<br>11.650368 | 14 | 19.1 | 1 |  | 3 x 10 meter<br>fyke nets + 6<br>cages |
| 10 | Tjolmenhamnen<br>Öckerö | 57.725139/<br>11.649242 | 14 | 17.4 | 1 | N = 2 | 3 x 10 meter<br>fyke nets + 6<br>cages |
| Ref 1 | Reference point 1<br>- west of<br>Långedrag | 57.665381/<br>11.816100 | 17 | 18.0 | 1 |  |  |
| Ref 2 | Reference point 2<br>- east of Vrångö | 57.574680/<br>11.805860 | 10 | 18.0 | 1 |  |  |
| (5.1) | Additional<br>Långedrag | 57.670277/<br>11.849265 | 17 | 16.0 | 1.9-2 |  |  |

**Table 4.**
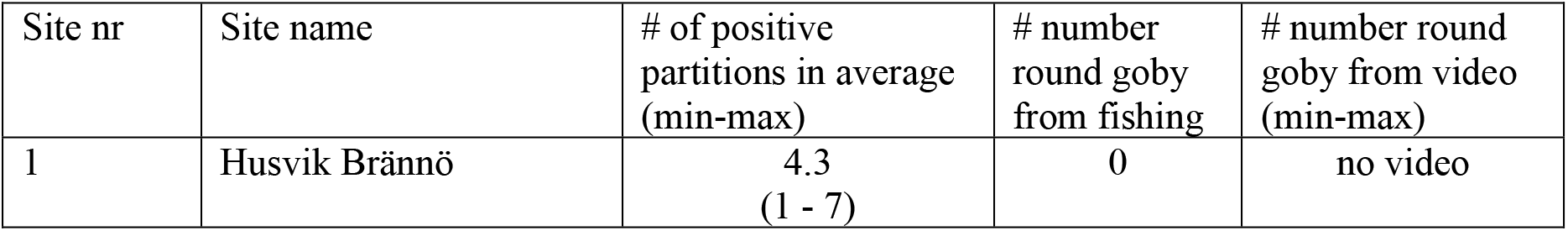
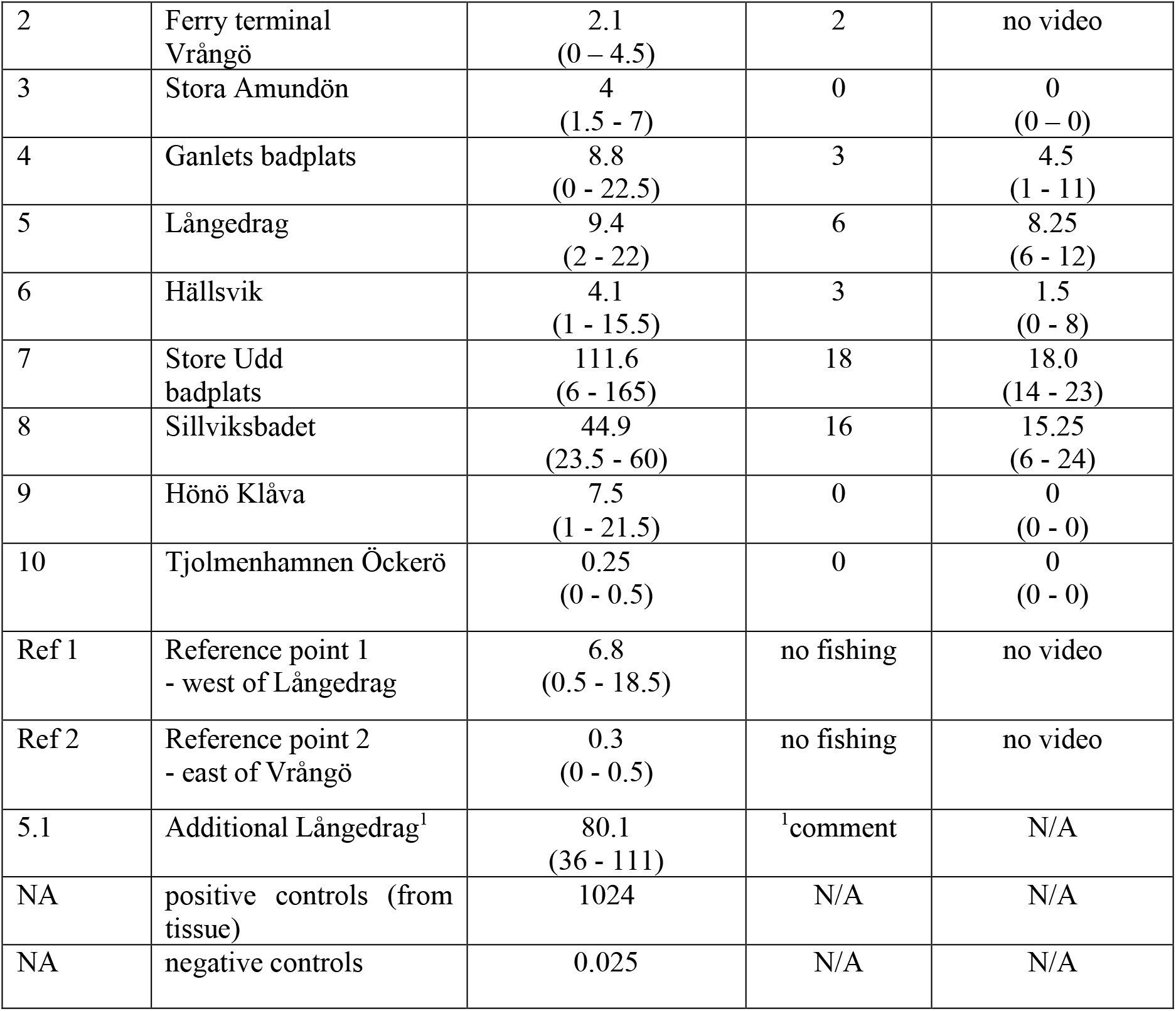
Numerical results from the eDNA analysis, fishing, and baited remote underwater video.

| Site nr | Site name | # of positive partitions in average (min-max) | # number round goby from fishing | # number round goby from video (min-max) |
| --- | --- | --- | --- | --- |
| 1 | Husvik Brännö | 4.3<br>(1 - 7) | 0 | no video |
| 2 | Ferry terminal<br>Vrångö | 2.1<br>(0 – 4.5) | 2 | no video |
| 3 | Stora Amundön | 4<br>(1.5 - 7) | 0 | 0<br>(0 – 0) |
| 4 | Ganlets badplats | 8.8<br>(0 - 22.5) | 3 | 4.5<br>(1 - 11) |
| 5 | Långedrag | 9.4<br>(2 - 22) | 6 | 8.25<br>(6 - 12) |
| 6 | Hällsvik | 4.1<br>(1 - 15.5) | 3 | 1.5<br>(0 - 8) |
| 7 | Store Udd<br>badplats | 111.6<br>(6 - 165) | 18 | 18.0<br>(14 - 23) |
| 8 | Sillviksbadet | 44.9<br>(23.5 - 60) | 16 | 15.25<br>(6 - 24) |
| 9 | Hönö Klåva | 7.5<br>(1 - 21.5) | 0 | 0<br>(0 - 0) |
| 10 | Tjolmenhamnen Öckerö | 0.25<br>(0 - 0.5) | 0 | 0<br>(0 - 0) |
| Ref 1 | Reference point 1<br>- west of Långedrag | 6.8<br>(0.5 - 18.5) | no fishing | no video |
| Ref 2 | Reference point 2<br>- east of Vrångö | 0.3<br>(0 - 0.5) | no fishing | no video |
| 5.1 | Additional Långedrag<sup>1</sup> | 80.1<br>(36 - 111) | <sup>1</sup>comment | N/A |
| NA | positive controls (from<br>tissue) | 1024 | N/A | N/A |
| NA | negative controls | 0.025 | N/A | N/A |

Extraction of DNA and subsequent analysis follows the protocol and scheme outlined in section 3.2 above.

### 3.5 Sampling by Baited Remote Underwater Video

At 8 of the 10 sites chosen for fishing and eDNA sampling, observational monitoring was conducted using baited remote underwater video systems (BRUVs) between the 13th to the 26th of September in 2021. These BRUVs were constructed from a 70x70 cm pvc frame, fitted with a camera (GoPro Hero 9, GoPro Inc., San Mateo, CA, USA) filming at 60 fps 2.4k and positioned at a 60 degree angle, 80 cm from a centered transparent plastic bait box (12 by 8 by 9 cm, fitted with holes for odor diffusion) baited with 70±10 gram of frozen-thawed shrimp (*Pandalus borealis*), previously known to attract *N. melanostomus* in fishing efforts by hook and line. These BRUVs were deployed by snorkeling, 2 at a time at a maximum depth of 2.5 meters, always angled with the camera pointing north for a similar angle of light, and both separated by at least 30 meters. Each site was filmed for one hour in two separate sessions, one in the morning and one in the afternoon, with less than 48 hours between each session resulting in 4 videos per site (except for site 10). The exact positioning for each BRUV was changed between sessions. Each video was then analyzed manually, and the maximum number of individuals > 10 cm total length observed at any one time during a film (Max N) was counted for *N. melanostomus*.

### 3.6 Sampling by fishing

See supplementary material for details of fishing methods. At ten of the field sites, fishing was conducted once between the 13^th^ to the 16^th^ of September 2021 at a depth of 2-5 meters using three double-ended fyke nets and five complementary crab and shrimp cages. The fishing gear was deployed for an average of around 26 hours (min 22h 24 min; max 23h 26min) from one day to the next. In total 50 individuals of *N. melanostomus* were caught at 6 of the 10 sampled sites. The majority (N = 47) were caught using fyke nets, while some were also caught in the cages at 2 localities (N = 3).

### 3.7 Statistical analysis

All statistical analyses were done using R (versions 4.1.0 and 4.3.1, R Core Team, 2021). The data from the experimental decay of round goby DNA in water was fitted to an exponential model using the ‘nls’ function in the drc package (Gerhard, Baty, Streibig, & D., 2015). The model is presented in section 4.1.

The amount of DNA found in the samples from the seascape outside of Gothenburg harbour (number of positive partitions), together with data on fishing catches (N per CPU) and maximum number of adult fish recorded with BRUV film (MaxN) was tested against each other for correlative relationships. Since all data was replicated, but the replicates were not expected to influence each other (i.e. the first sample of eDNA on a date was not expected to specifically correlate to the first video taken on the same site another date), all replicates were pooled by site and an average calculated to use for the correlations. The three possible correlations (average eDNA ∼ average recorded MaxN; average eDNA ∼ N per CPU; average recorded MaxN ∼ N per CPU) were first modelled using the ‘lm’ function in the packages lme4 (Bates et al., 2015) and car (Fox et al., 2013). The models were explored for normality by visually inspecting the residuals vs fitted values, the frequency distribution of residuals, the theoretical and observed quantiles and high influence points, using the ‘plot(model)’ function. Since the data was not normally distributed, a non-parametric Spearman’s rank correlation test was then used to model the relationships between the parameters using the package stats (R Core Team, 2023) and presented below in section 4.2.

## 4.0 Results

### 4.1 eDNA decay rate

The results of the eDNA decaying rate indicate that the amount of potentially detectable DNA from a target source decreases rather rapidly (Figure 1). The half-life in this experiment is estimated to be around 10 hours at 15 PSU salinity and 15°C, which is consistent with results in other studies of aquatic metazoans (Andruszkiewicz Allan et al. 2020).

**Figure 1:**
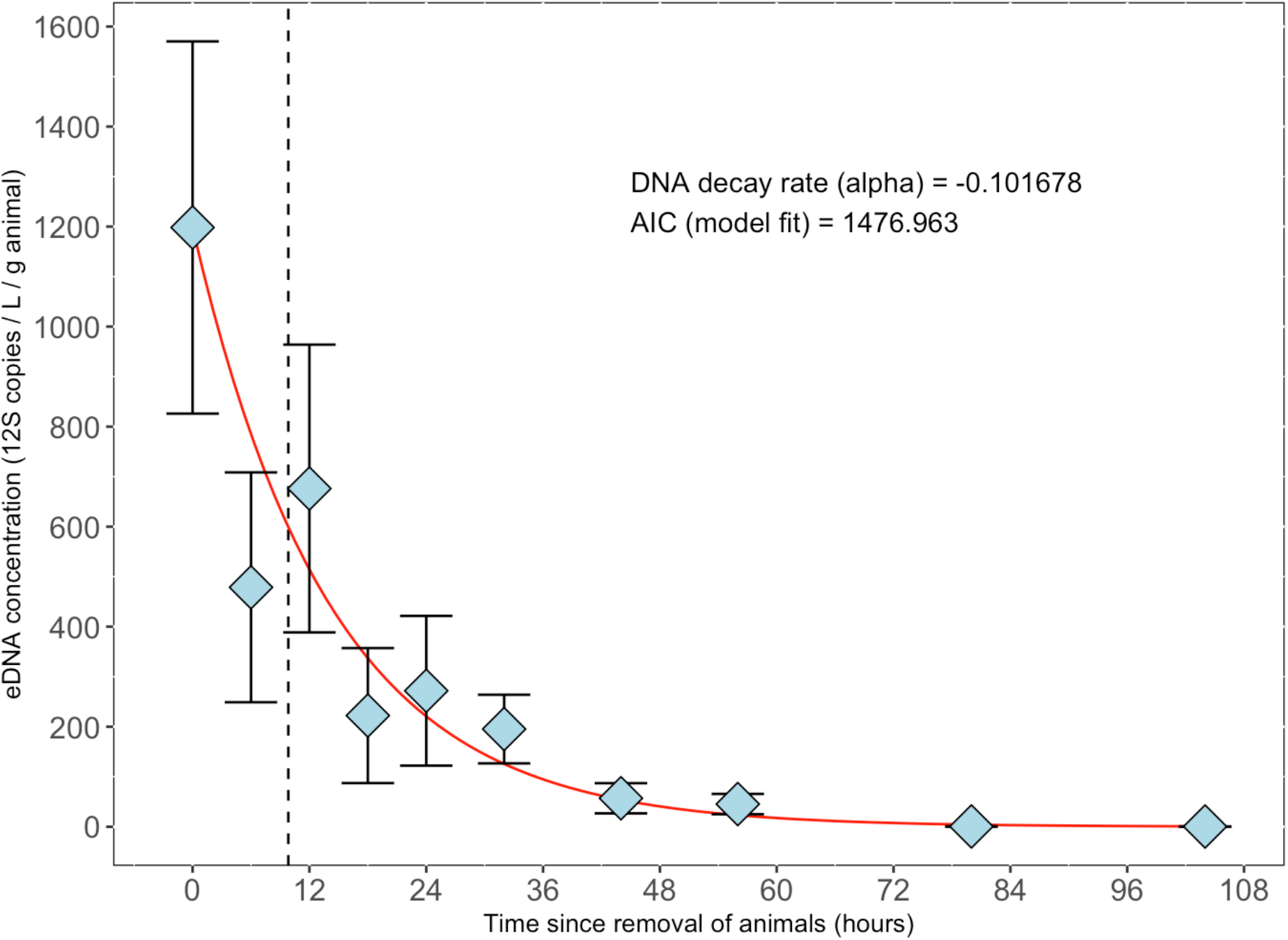
Exponential decay of eDNA from *Neogobius melanostomus* after removal from 30 liter tanks where the animals were kept during 24 hours. Diamonds show the mean number of positive partitions (eDNA copies) / Liter water / gram animal. Error bars show standard error. Red line shows decay model fitted using nls function. Vertical dashed line shows half-life (9.8 hours).

### 4.2 eDNA verified and compared to fishing efforts and BRUV observations

Sampling was carried out in shallow, and sheltered, water just inside the jetty east of Site 5 “Långedrag”. Although no simultaneous fishing was done at the moment of water sampling, later line hooking of round goby for another experiment caught eight fish in just an hour indicating that the species is abundant at this site.

### 4.3 Correlation between eDNA copy number and fish abundance

Significant correlative relationships were found between the mean number of eDNA copies and the mean number of maximum adult round gobies recorded at the 8 sites where both parameters were sampled (Spearman’s rank correlation, R_s_ = 0.854, S = 12.271, p = 0.007) (Figure 2). A significant correlative relationship was also found between the mean number of eDNA copies the number of fish caught at the ten sites where eDNA was also sampled (Spearman’s rank correlation, R_s_ = 0.719, S = 46.348, p-value = 0.019). Furthermore, an almost linear relationship was found between the mean number of maximum adult round gobies recorded and the number of fish caught per site (Spearman’s rank correlation, R_s_ = 1, S = 0, p = < 0.001). However, among the sites sampled, only one was found to lack any round goby eDNA signal in any sample taken (Site 10) and completely lack round goby recorded on BRUV or caught by fishing. In contrast, at some sites (Sites 2, 3 and 9), round goby eDNA was found (albeit at low levels) but no round gobies were recorded on the four BRUVs or caught in any of the gear placed in the area. Furthermore, samples from Site 4 and Site 6 showed a very small eDNA signal (some with 0 copies found) despite these sites having up to 12 round gobies on camera and also a few caught by fishing. Although the correlations with the number of eDNA positive partitions from the data at hand shows a clear connection with fish population abundance (like e.g. Sites 7 and 8), one should bear in mind that there are many environmental factors influencing the sampling and how many DNA molecules can be retrieved - see also below in “Discussion”.

**Figure 2.**
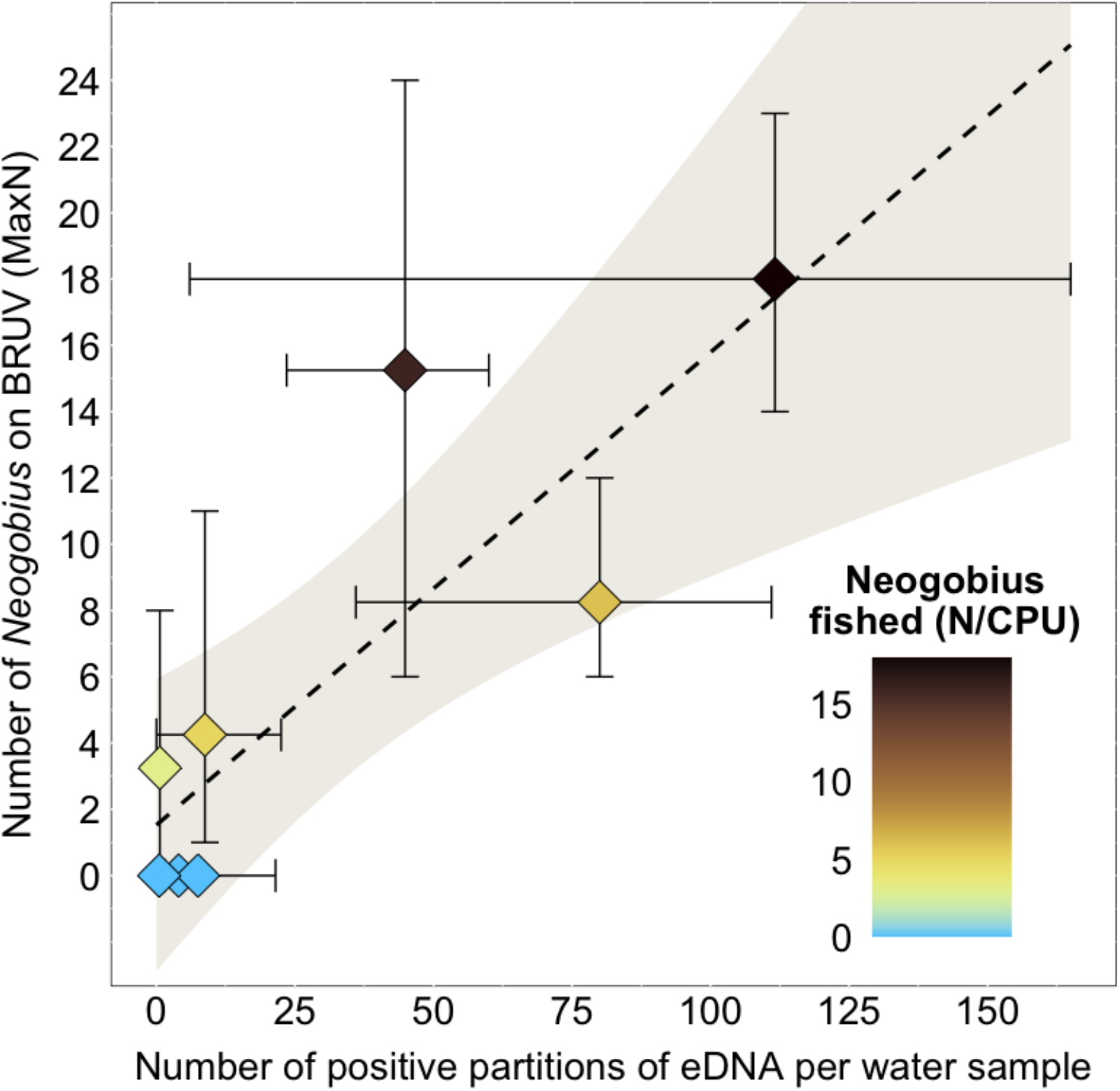
Correlations of eDNA copy numbers (positive partitions per water sample) and the number of adult round gobies recorded and caught at each site. Y-axis show the mean value of the maximum number of round gobies shown during one time on camera from four samples taken with baited remote underwater video (BRUV) and the mean value of eDNA copies from four samples of water. Color of diamonds show the number of fish caught per unit effort (N/CPU) per site. Error bars show min and max values from the four samples taken from each site. Regression line shows a fitted linear model to the mean values, shaded area shows 95% confidence interval.

## 5.0 Discussion

In this study, we developed a target species dPCR assay for round goby (*Neogobius melanostomus*) and verified its applicability to distinguish the species from other common gobiids in our study area. We then experimentally determined the decaying rate of round goby DNA in water to a half-life of 9.8 hours in 15 PSU and 15°C. Finally, we sampled 10 sites within a 400 km^2^ area for eDNA and assessed presence of the species using fyke-nets and baited remote video to validate the accuracy of the water samples to predict presence, and abundance. We found that the number of DNA copies extracted from the water samples varied strongly at sites where round gobies were caught in nets or on video, but that the average value from four water samples significantly correlated with an average value from four video samples, and also with the total catch at each site. These results show that it is possible to predict the presence of this invasive species in an area using eDNA, and with sufficient amounts of replicates, potentially determine the relative abundance between multiple sites.

In our experiment, the results show a clear decay of the target DNA fragment over time. The half-life of 9.8 hours in 15 PSU and 15°C, and without direct sunlight, shows that even in conditions that are stable, with little biological interference from microorganisms (artificial seawater was used for housing the fish), rapid decaying of DNA does take place. Previous experimental studies have shown half-life rates are taxonomic group and species specific (Allan et al., 2020). Among teleosts, DNA copy half-lifes vary in the magnitude of hours to days, with hours being more common (Allan et al., 2020). For example: between 6.8 and 12.6 hours (for five pacific species of anchovies, sardines and mackerel) at 18.7-22.0°C and 31.1-39.2 PSU salinity (Sassoubre et al., 2016); 1.22-3.21 hours for winter flounder, 2.89-9.9 hours for summer flounder and 3.71-4.0 hours for black sea bass at 15.9-19.9°C and 27.9-33.9 PSU salinity (Kirtane et al., 2021). Muskellunge in freshwater has shown half-lifes for eDNA from larvae at 2.7-3.7 hours at 15°C, and from juveniles at 5.2-10.8 hours at 20°C (Wilder et al., 2023). The round goby DNA decay rates, experimentally tested in stable salinity and with little bacterial load, is likely to be faster in a natural environment. This is due to that DNA half-life decreases with microbial load (Lance et al., 2017) and temperature (Lance et al., 2017; Allan et al., 2020) as well as from interactions with other particles (Harrison et al., 2019). Since eDNA often binds to particles and sinks (Hansen et al., 2018), our reported decay rates together with the high site fidelity of round gobies (Kornis et al., 2012), makes it highly likely that our samples are representative of the species presence at the time of sampling. All samples for this study were taken within the known distribution range of round goby on the Swedish west coast, and positive samples of eDNA showed up at every single site (but not in all samples). Among the sites where no round goby presence could be validated by the other methods (i.e. caught by fishing or seen in video, site 1, 3, 9 and 10), records of them exist from sources outside of this study. At site 1, no fish were found during video sampling in 2022, but 2 individuals have been caught within 1200 meters by citizen scientists in 2018 and 2019 respectively (Swedish species observation system, www.artportalen.se, accessed 25th of January 2024). At site 3 (1.5-7 partitions detected) and 10 (0-0.5 partitions detected), round gobies were found in low numbers (N = 1) when video-sampling using similar methodology in 2022 (4 BRUVs, 1 h each), the year after this study was conducted (Green et al., unpublished). At site 9 (1-21.5 partitions detected), video sampling was conducted in 2022, and in 2023 with the same effort as for site 3 and 10 above, and two individuals (one male and one female) were observed on the same video in 2023. This shows that all sites where eDNA was found in the water samples also very likely had a presence of the species and that its unlikely that false positives were encountered.

A limited understanding of the fate and dynamics of eDNA in water, at the time of writing, is likely the largest barrier to abundance level measurements (Wang et al., 2021). Though the aim of this study was not to test what local factors contribute to the stochasticity of the amount of DNA found in the water sample, the ones we expect to have the largest effect on DNA transport, retention and resuspension, apart from the species’ density are listed here. Firstly, the round goby is a benthic fish that spends most of its time on the bottom (Kornis et al., 2012), though it also positions itself on walls and other structures (Bussmann et al., 2022). It utilizes crevices and holes for breeding and shelter (Kornis et al., 2012) and migrates to deeper water during winter (Behrens et al., 2022). Round gobies are also known to be worse swimmers than similarly sized and shaped fishes (Wiegleb et al., 2022) and they generally avoid water with high velocities (Roje et al., 2021). The shedding of the DNA is therefore expected to occur close to the bottom sediment, close to walls and in crevices and holes, and importantly - not in the pelagic/water column. Second, the biota on and around these structures are also likely to influence the presence of DNA in the water, the most important effect likely coming from filter-feeding invertebrates (such as bivalves), that rapidly clear the water from particles, along with any DNA attached to them (Friebertshauser et al., 2018). Round gobies are known to predate on bivalves (Foley et al., 2017, Karlsson et al., 2007) and are expected to occur in areas with high bivalve abundance (shallow coastal zones with high chlorophyll content and wave action - Holmes et al., 2019) so accounting for their presence will be ecologically relevant. Third, experimental studies have also shown that round gobies prefer to spend time in shelter during daytime (Roje et al., 2021), and instead spend time in calm, quiet waters with low turbulence during the hours of darkness (Roje et al., 2021). Time of day will likely have a large effect on the amount of eDNA captured in a sample (Reinholdt Jensen et al., 2022). Fourth, local conditions at a site, due to weather and the physical attributes of the area (sheltered zones, differences in depth) will likely have a strong influence on the quantity of DNA in the water sample. Waves and currents mix water more rapidly, likely affecting resuspension and transport to a large degree (Harrison et al., 2019). Shedding rates have been shown to increase with density in experimental conditions (Lance et al., 2017; Minegishi et al. 2019). In summary, samples are expected to be more representative within close proximity to the bottom, and in calmer conditions, when the fish are more active such as during the night or early morning. At exposed sites, more prone to stronger current conditions, one should sample from leeward positions if possible. The sampling from the two sites at Långedrag exemplifies this situation. The first sample was taken outside the pier (site 5) under windy conditions and more currents, while the second site (“additional Långedrag”) was from the sheltered inside of the same pier.

## 6.0 Conclusions

In this study, we show that eDNA can be used to verify the current presence, and to a degree, assess the relative abundance of the invasive round goby *Neogobius melanostomus* in a marine seascape. Our findings contribute to the understanding of eDNA characteristics and dynamics in the marine environment and may help us to interpret future results of eDNA surveillance.

## 7.0 Acknowledgements

Jannic Wochen conducted a substantial part of the laboratory work during his contract with SeAnalytics. Jimmy Ahlsén and Johanna Bergkvist at Marine Monitoring were contracted to support the project with the fishing survey. We are very grateful to Christin Appelqvist for her help with obtaining gobiid samples and to Charlotta Kvarnemo for the help with the gobiid species identification. A part of the lab work (DNA extraction and conventional PCR tests) was conducted by students Diana Checkmous and Hanna Basic from Strömstad high school, as a high school research project under the direct supervision of Marina Panova, and we thank Diana and Hanna for their great work in the lab. The study was financially supported by the Swedish Environmental Protection Agency and Swedish Agency for Marine and Water Management (contract 802-0107-18) to Per Sundberg and (grant nr 2020-00055) to Leon Green.

## 9.0 Data availability statement

Raw data supporting this study will be available following submission and publication in a peer-review journal.

## References

Adrian-Kalchhauser, I., & Burkhardt-Holm, P. (2016). An eDNA Assay to Monitor a Globally Invasive Fish Species from Flowing Freshwater. PLOS ONE, 11(1), e0147558. 10.1371/journal.pone.0147558

Ahmed, D. A., Hudgins, E. J., Cuthbert, R. N., Kourantidou, M., Diagne, C., Haubrock, P. J., Leung, B., Liu, C., Leroy, B., Petrovskii, S., Beidas, A., & Courchamp, F. (2022). Managing biological invasions: The cost of inaction. Biological Invasions, 24(7), 1927–1946. 10.1007/s10530-022-02755-0

Andruszkiewicz Allan, E., Zhang, W. G., C. Lavery, A., & F. Govindarajan, A. (2021). Environmental DNA shedding and decay rates from diverse animal forms and thermal regimes. Environmental DNA, 3(2), 492–514. 10.1002/edn3.141

Anton, A., Geraldi, N. R., Lovelock, C. E., Apostolaki, E. T., Bennett, S., Cebrian, J., Krause-Jensen, D., Marbà, N., Martinetto, P., Pandolfi, J. M., Santana-Garcon, J., & Duarte, C. M. (2019). Global ecological impacts of marine exotic species. Nature Ecology & Evolution, 3(5), 787–800. 10.1038/s41559-019-0851-0

Bates, D., Mächler, M., Bolker, B., & Walker, S. (2015). Fitting Linear Mixed-Effects Models Using lme4. Journal of Statistical Software, 67(1). 10.18637/jss.v067.i01

Bax, N., Carlton, J. T., Mathews-Amos, A., Haedrich, R. L., Howarth, F. G., Purcell, J. E., Rieser, A., & Gray, A. (2001). The Control of Biological Invasions in the World’s Oceans. Conservation Biology, 15(5), 1234–1246. 10.1111/j.1523-1739.2001.99487.x

Behrens, J. W., Ryberg, M. P., Einberg, H., Eschbaum, R., Florin, A.-B., Grygiel, W., Herrmann, J. P., Huwer, B., Hüssy, K., Knospina, E., Nõomaa, K., Oesterwind, D., Polte, P., Smoliński, S., Ustups, D., Van Deurs, M., & Ojaveer, H. (2022). Seasonal depth distribution and thermal experience of the non-indigenous round goby Neogobius melanostomus in the Baltic Sea: Implications to key trophic relations. Biological Invasions, 24(2), 527–541. 10.1007/s10530-021-02662-w

Beng, K. C., & Corlett, R. T. (2020). Applications of environmental DNA (eDNA) in ecology and conservation: Opportunities, challenges and prospects. Biodiversity and Conservation, 29(7), 2089–2121. 10.1007/s10531-020-01980-0

Bio-Rad. (n.d.). Droplet DigitalTM PCR Applications Guide (Bulletin 6407 Ver B) [Application guide]. Retrieved May 2, 2024, from https://www.bio-rad.com/webroot/web/pdf/lsr/literature/Bulletin_6407.pdf

Bonebrake, T. C., Brown, C. J., Bell, J. D., Blanchard, J. L., Chauvenet, A., Champion, C., Chen, I., Clark, T. D., Colwell, R. K., Danielsen, F., Dell, A. I., Donelson, J. M., Evengård, B., Ferrier, S., Frusher, S., Garcia, R. A., Griffis, R. B., Hobday, A. J., Jarzyna, M. A., … Pecl, G. T. (2018). Managing consequences of climate-driven species redistribution requires integration of ecology, conservation and social science. Biological Reviews, 93(1), 284–305. 10.1111/brv.12344

Bradley, B. A., Laginhas, B. B., Whitlock, R., Allen, J. M., Bates, A. E., Bernatchez, G., Diez, J. M., Early, R., Lenoir, J., Vilà, M., & Sorte, C. J. B. (2019). Disentangling the abundance–impact relationship for invasive species. Proceedings of the National Academy of Sciences, 116(20), 9919–9924. 10.1073/pnas.1818081116

Bussmann, K., Hirsch, P. E., Lehmann, M. F., & Burkhardt-Holm, P. (2022). Differential habitat use of a notorious invasive fish, the round goby, in a translocation-relevant system. Ecology and Evolution, 12(8), e9202. 10.1002/ece3.9202

Collins, R. A., Wangensteen, O. S., O’Gorman, E. J., Mariani, S., Sims, D. W., & Genner, M. J. (2018). Persistence of environmental DNA in marine systems. Communications Biology, 1(1), 185. 10.1038/s42003-018-0192-6

Cuthbert, R. N., Diagne, C., Hudgins, E. J., Turbelin, A., Ahmed, D. A., Albert, C., Bodey, T. W., Briski, E., Essl, F., Haubrock, P. J., Gozlan, R. E., Kirichenko, N., Kourantidou, M., Kramer, A. M., & Courchamp, F. (2022). Biological invasion costs reveal insufficient proactive management worldwide. Science of The Total Environment, 819, 153404. 10.1016/j.scitotenv.2022.153404

Cuthbert, R. N., Pattison, Z., Taylor, N. G., Verbrugge, L., Diagne, C., Ahmed, D. A., Leroy, B., Angulo, E., Briski, E., Capinha, C., Catford, J. A., Dalu, T., Essl, F., Gozlan, R. E., Haubrock, P. J., Kourantidou, M., Kramer, A. M., Renault, D., Wasserman, R. J., & Courchamp, F. (2021). Global economic costs of aquatic invasive alien species. Science of The Total Environment, 775, 145238. 10.1016/j.scitotenv.2021.145238

Danziger, A. M., & Frederich, M. (2022). Challenges in eDNA detection of the invasive European green crab, Carcinus maenas. Biological Invasions, 24(6), 1881–1894. 10.1007/s10530-022-02757-y

Deiner, K., & Altermatt, F. (2014). Transport Distance of Invertebrate Environmental DNA in a Natural River. PLoS ONE, 9(2), e88786. 10.1371/journal.pone.0088786

Doi, H., Takahara, T., Minamoto, T., Matsuhashi, S., Uchii, K., & Yamanaka, H. (2015). Droplet Digital Polymerase Chain Reaction (PCR) Outperforms Real-Time PCR in the Detection of Environmental DNA from an Invasive Fish Species. Environmental Science & Technology, 49(9), 5601–5608. 10.1021/acs.est.5b00253

Doyle, J., & Uthicke, S. (2021). Sensitive environmental DNA detection via lateral flow assay (dipstick)—A case study on corallivorous crown-of-thorns sea star (Acanthaster cf. Solaris) detection. Environmental DNA, 3(2), 323–342. 10.1002/edn3.123

Early, R., Bradley, B. A., Dukes, J. S., Lawler, J. J., Olden, J. D., Blumenthal, D. M., Gonzalez, P., Grosholz, E. D., Ibañez, I., Miller, L. P., Sorte, C. J. B., & Tatem, A. J. (2016). Global threats from invasive alien species in the twenty-first century and national response capacities. Nature Communications, 7(1), 12485. 10.1038/ncomms12485

Foley, C. J., Andree, S. R., Pothoven, S. A., Nalepa, T. F., & Höök, T. O. (2017). Quantifying the predatory effect of round goby on Saginaw Bay dreissenids. Journal of Great Lakes Research, 43(1), 121–131. 10.1016/j.jglr.2016.10.018

Friebertshauser, R., Shollenberger, K., Janosik, A., Garner, J. T., & Johnston, C. (2019). The effect of bivalve filtration on eDNA-based detection of aquatic organisms. PLOS ONE, 14(11), e0222830. 10.1371/journal.pone.0222830

Giménez, L., Exton, M., Spitzner, F., Meth, R., Ecker, U., Jungblut, S., Harzsch, S., Saborowski, R., & Torres, G. (2020). Exploring larval phenology as predictor for range expansion in an invasive species. Ecography, 43(10), 1423–1434. 10.1111/ecog.04725

Gold, Z., Wall, A. R., Schweizer, T. M., Pentcheff, N. D., Curd, E. E., Barber, P. H., Meyer, R. S., Wayne, R., Stolzenbach, K., Prickett, K., Luedy, J., & Wetzer, R. (2022). A manager’s guide to using eDNA metabarcoding in marine ecosystems. PeerJ, 10, e14071. 10.7717/peerj.14071

Green, L., Apostolou, A., Faust, E., Palmqvist, K., Behrens, J. W., Havenhand, J. N., Leder, E. H., & Kvarnemo, C. (2021). Ancestral Sperm Ecotypes Reveal Multiple Invasions of a Non-Native Fish in Northern Europe. Cells, 10(7), 1743. 10.3390/cells10071743

Green, L., Faust, E., Hinchcliffe, J., Brijs, J., Holmes, A., Englund Örn, F., Svensson, O., Roques, J. A. C., Leder, E. H., Sandblom, E., & Kvarnemo, C. (2023). Invader at the edge—Genomic origins and physiological differences of round gobies across a steep urban salinity gradient. Evolutionary Applications, 16(2), 321–337. 10.1111/eva.13437

Green, S. J., & Grosholz, E. D. (2021). Functional eradication as a framework for invasive species control. Frontiers in Ecology and the Environment, 19(2), 98–107. 10.1002/fee.2277

Hansen, B. K., Bekkevold, D., Clausen, L. W., & Nielsen, E. E. (2018). The sceptical optimist: Challenges and perspectives for the application of environmental DNA in marine fisheries. Fish and Fisheries, 19(5), 751–768. 10.1111/faf.12286

Harrison, J. B., Sunday, J. M., & Rogers, S. M. (2019). Predicting the fate of eDNA in the environment and implications for studying biodiversity. Proceedings of the Royal Society B: Biological Sciences, 286(1915), 20191409. 10.1098/rspb.2019.1409

Haubrock, P. J., Bernery, C., Cuthbert, R. N., Liu, C., Kourantidou, M., Leroy, B., Turbelin, A. J., Kramer, A. M., Verbrugge, L. N. H., Diagne, C., Courchamp, F., & Gozlan, R. E. (2022). Knowledge gaps in economic costs of invasive alien fish worldwide. Science of The Total Environment, 803, 149875. 10.1016/j.scitotenv.2021.149875

Holman, L. E., De Bruyn, M., Creer, S., Carvalho, G., Robidart, J., & Rius, M. (2019). Detection of introduced and resident marine species using environmental DNA metabarcoding of sediment and water. Scientific Reports, 9(1), 11559. 10.1038/s41598-019-47899-7

Holmes, M., Kotta, J., Persson, A., & Sahlin, U. (2019). Marine protected areas modulate habitat suitability of the invasive round goby (Neogobius melanostomus) in the Baltic Sea. Estuarine, Coastal and Shelf Science, 229, 106380. 10.1016/j.ecss.2019.106380

Hunter, M. E., Dorazio, R. M., Butterfield, J. S. S., Meigs-Friend, G., Nico, L. G., & Ferrante, J. A. (2017). Detection limits of quantitative and digital PCR assays and their influence in presence– absence surveys of environmental DNA. Molecular Ecology Resources, 17(2), 221–229. 10.1111/1755-0998.12619

ICES. (2022). Workshop on stickleback and round goby in the Baltic Sea (WKSTARGATE) (p. 2298726 Bytes). ICES Scientific Reports. 10.17895/ICES.PUB.21345291.V1

Jensen, M. R., Sigsgaard, E. E., Ávila, M. D. P., Agersnap, S., Brenner-Larsen, W., Sengupta, M. E., Xing, Y., Krag, M. A., Knudsen, S. W., Carl, H., Møller, P. R., & Thomsen, P. F. (2022). Short-term temporal variation of coastal marine eDNA. Environmental DNA, 4(4), 747–762. 10.1002/edn3.285

Karlson, A. M. L., Almqvist, G., Skóra, K. E., & Appelberg, M. (2007). Indications of competition between non-indigenous round goby and native flounder in the Baltic Sea. ICES Journal of Marine Science, 64(3), 479–486. 10.1093/icesjms/fsl049

Kirtane, A., Wieczorek, D., Noji, T., Baskin, L., Ober, C., Plosica, R., Chenoweth, A., Lynch, K., & Sassoubre, L. (2021). Quantification of Environmental DNA (eDNA) shedding and decay rates for three commercially harvested fish species and comparison between eDNA detection and trawl catches. Environmental DNA, 3(6), 1142–1155. 10.1002/edn3.236

Knudsen, S. W., Hesselsøe, M., Thaulow, J., Agersnap, S., Hansen, B. K., Jacobsen, M. W., Bekkevold, D., Jensen, S. K. S., Møller, P. R., & Andersen, J. H. (2022). Monitoring of environmental DNA from nonindigenous species of algae, dinoflagellates and animals in the North East Atlantic. Science of The Total Environment, 821, 153093. 10.1016/j.scitotenv.2022.153093

Kornis, M. S., Mercado-Silva, N., & Vander Zanden, M. J. (2012). Twenty years of invasion: A review of round goby Neogobius melanostomus biology, spread and ecological implications. Journal of Fish Biology, 80(2), 235–285. 10.1111/j.1095-8649.2011.03157.x

Lance, R., Klymus, K., Richter, C., Guan, X., Farrington, H., Carr, M., Thompson, N., Chapman, D., & Baerwaldt, K. (2017). Experimental observations on the decay of environmental DNA from bighead and silver carps. Management of Biological Invasions, 8(3), 343–359. 10.3391/mbi.2017.8.3.08

Maruyama, A., Nakamura, K., Yamanaka, H., Kondoh, M., & Minamoto, T. (2014). The Release Rate of Environmental DNA from Juvenile and Adult Fish. PLoS ONE, 9(12), e114639. 10.1371/journal.pone.0114639

McConnachie, M. M., Van Wilgen, B. W., Ferraro, P. J., Forsyth, A. T., Richardson, D. M., Gaertner, M., & Cowling, R. M. (2016). Using counterfactuals to evaluate the cost-effectiveness of controlling biological invasions. Ecological Applications, 26(2), 475–483. 10.1890/15-0351

Miller, A. W., & Ruiz, G. M. (2014). Arctic shipping and marine invaders. Nature Climate Change, 4(6), 413–416. 10.1038/nclimate2244

Minegishi, Y., Wong, M. K.-S., Kanbe, T., Araki, H., Kashiwabara, T., Ijichi, M., Kogure, K., & Hyodo, S. (2019). Spatiotemporal distribution of juvenile chum salmon in Otsuchi Bay, Iwate, Japan, inferred from environmental DNA. PLOS ONE, 14(9), e0222052. 10.1371/journal.pone.0222052

Miya, M. (2022). Environmental DNA Metabarcoding: A Novel Method for Biodiversity Monitoring of Marine Fish Communities. Annual Review of Marine Science, 14(1), 161–185. 10.1146/annurev-marine-041421-082251

Miya, M., Sato, Y., Fukunaga, T., Sado, T., Poulsen, J. Y., Sato, K., Minamoto, T., Yamamoto, S., Yamanaka, H., Araki, H., Kondoh, M., & Iwasaki, W. (2015). MiFish, a set of universal PCR primers for metabarcoding environmental DNA from fishes: Detection of more than 230 subtropical marine species. Royal Society Open Science, 2(7), 150088. 10.1098/rsos.150088

Morin, F., Panova, M. A. Z., Schweizer, M., Wiechmann, M., Eliassen, N., Sundberg, P., Cluzel-Burgalat, L., & Polovodova Asteman, I. (2023). Hidden aliens: Application of digital PCR to track an exotic foraminifer across the Skagerrak (North Sea) correlates well with traditional morphospecies analysis. Environmental Microbiology, 25(11), 2321–2337. 10.1111/1462-2920.16458

Nygård, H., Oinonen, S., Hällfors, H. A., Lehtiniemi, M., Rantajärvi, E., & Uusitalo, L. (2016). Price vs. Value of Marine Monitoring. Frontiers in Marine Science, 3. 10.3389/fmars.2016.00205

Pyšek, P., Hulme, P. E., Simberloff, D., Bacher, S., Blackburn, T. M., Carlton, J. T., Dawson, W., Essl, F., Foxcroft, L. C., Genovesi, P., Jeschke, J. M., Kühn, I., Liebhold, A. M., Mandrak, N. E., Meyerson, L. A., Pauchard, A., Pergl, J., Roy, H. E., Seebens, H., … Richardson, D. M. (2020). Scientists’ warning on invasive alien species. Biological Reviews, 95(6), 1511–1534. 10.1111/brv.12627

Rahel, F. J., & Olden, J. D. (2008). Assessing the Effects of Climate Change on Aquatic Invasive Species. Conservation Biology, 22(3), 521–533. 10.1111/j.1523-1739.2008.00950.x

Robertson, P. A., Mill, A., Novoa, A., Jeschke, J. M., Essl, F., Gallardo, B., Geist, J., Jarić, I., Lambin, X., Musseau, C., Pergl, J., Pyšek, P., Rabitsch, W., Von Schmalensee, M., Shirley, M., Strayer, D. L., Stefansson, R. A., Smith, K., & Booy, O. (2020). A proposed unified framework to describe the management of biological invasions. Biological Invasions, 22(9), 2633–2645. 10.1007/s10530-020-02298-2

Roje, S., Drozd, B., Richter, L., Kubec, J., Polívka, Z., Worischka, S., & Buřič, M. (2021). Comparison of Behavior and Space Use of the European Bullhead Cottus gobio and the Round Goby Neogobius melanostomus in a Simulated Natural Habitat. Biology, 10(9), 821. 10.3390/biology10090821

Rourke, M. L., Fowler, A. M., Hughes, J. M., Broadhurst, M. K., DiBattista, J. D., Fielder, S., Wilkes Walburn, J., & Furlan, E. M. (2022). Environmental DNA (eDNA) as a tool for assessing fish biomass: A review of approaches and future considerations for resource surveys. Environmental DNA, 4(1), 9–33. 10.1002/edn3.185

Roy, H. E., Pauchard, A., Stoett, P., Renard Truong, T., Bacher, S., Galil, B. S., Hulme, P. E., Ikeda, T., Sankaran, K., McGeoch, M. A., Meyerson, L. A., Nuñez, M. A., Ordonez, A., Rahlao, S. J., Schwindt, E., Seebens, H., Sheppard, A. W., & Vandvik, V. (2024). IPBES Invasive Alien Species Assessment: Summary for Policymakers (Version 3). [object Object]. 10.5281/ZENODO.7430692

Sardain, A., Sardain, E., & Leung, B. (2019). Global forecasts of shipping traffic and biological invasions to 2050. Nature Sustainability, 2(4), 274–282. 10.1038/s41893-019-0245-y

Sassoubre, L. M., Yamahara, K. M., Gardner, L. D., Block, B. A., & Boehm, A. B. (2016). Quantification of Environmental DNA (eDNA) Shedding and Decay Rates for Three Marine Fish. Environmental Science & Technology, 50(19), 10456–10464. 10.1021/acs.est.6b03114

Sepulveda, A. J., Nelson, N. M., Jerde, C. L., & Luikart, G. (2020). Are Environmental DNA Methods Ready for Aquatic Invasive Species Management? Trends in Ecology & Evolution, 35(8), 668–678. 10.1016/j.tree.2020.03.011

Shaw, J. L. A., Clarke, L. J., Wedderburn, S. D., Barnes, T. C., Weyrich, L. S., & Cooper, A. (2016). Comparison of environmental DNA metabarcoding and conventional fish survey methods in a river system. Biological Conservation, 197, 131–138. 10.1016/j.biocon.2016.03.010

Shogren, A. J., Tank, J. L., Andruszkiewicz, E., Olds, B., Mahon, A. R., Jerde, C. L., & Bolster, D. (2017). Controls on eDNA movement in streams: Transport, Retention, and Resuspension. Scientific Reports, 7(1), 5065. 10.1038/s41598-017-05223-1

Simberloff, D., Martin, J.-L., Genovesi, P., Maris, V., Wardle, D. A., Aronson, J., Courchamp, F., Galil, B., García-Berthou, E., Pascal, M., Pyšek, P., Sousa, R., Tabacchi, E., & Vilà, M. (2013). Impacts of biological invasions: What’s what and the way forward. Trends in Ecology & Evolution, 28(1), 58–66. 10.1016/j.tree.2012.07.013

Staehr, P. A. U., Dahl, K., Buur, H., Göke, C., Sapkota, R., Winding, A., Panova, M., Obst, M., & Sundberg, P. (2022). Environmental DNA Monitoring of Biodiversity Hotspots in Danish Marine Waters. Frontiers in Marine Science, 8, 800474. 10.3389/fmars.2021.800474

Stockwell, D. R. B., & Peterson, A. T. (2002). Effects of sample size on accuracy of species distribution models. Ecological Modelling, 148(1), 1–13. 10.1016/S0304-3800(01)00388-X

Sundberg, P., Axberg, A., Bravell, F., Wocken, Y., Ahlsen, J., Bergkvist, J., & Magnusson, M. (2022). Övervakning av svartmunnad smörbult—Pilotstudie med eDNA och provfiske i Göteborgs skärgård 2021 (Rapport 2022:14.; p. 26). Länsstyrelsen Västra Götaland. https://www.lansstyrelsen.se/vastra-gotaland/om-oss/vara-tjanster/publikationer/2022/overvakning-av-svartmunnad-smorbult---pilotstudie-med-edna-och-provfiske-i-goteborgs-skargard-2021.html

Thalinger, B., Deiner, K., Harper, L. R., Rees, H. C., Blackman, R. C., Sint, D., Traugott, M., Goldberg, C. S., & Bruce, K. (2021). A validation scale to determine the readiness of environmental DNA assays for routine species monitoring. Environmental DNA, 3(4), 823–836. 10.1002/edn3.189

Thomsen, P. F., Kielgast, J., Iversen, L. L., Møller, P. R., Rasmussen, M., & Willerslev, E. (2012). Detection of a Diverse Marine Fish Fauna Using Environmental DNA from Seawater Samples. PLoS ONE, 7(8), e41732. 10.1371/journal.pone.0041732

Thresher, R. E., & Kuris, A. M. (2004). Options for Managing Invasive Marine Species. Biological Invasions, 6(3), 295–300. 10.1023/B:BINV.0000034598.28718.2e

Uthicke, S., Lamare, M., & Doyle, J. R. (2018). eDNA detection of corallivorous seastar (Acanthaster cf. Solaris) outbreaks on the Great Barrier Reef using digital droplet PCR. Coral Reefs, 37(4), 1229–1239. 10.1007/s00338-018-1734-6

Van Deurs, M., Moran, N. P., Schreiber Plet-Hansen, K., Dinesen, G. E., Azour, F., Carl, H., Møller, P. R., & Behrens, J. W. (2021). Impacts of the invasive round goby (Neogobius melanostomus) on benthic invertebrate fauna: A case study from the Baltic Sea. NeoBiota, 68, 19–30. 10.3897/neobiota.68.67340

Wang, S., Yan, Z., Hänfling, B., Zheng, X., Wang, P., Fan, J., & Li, J. (2021). Methodology of fish eDNA and its applications in ecology and environment. Science of The Total Environment, 755, 142622. 10.1016/j.scitotenv.2020.142622

Wiegleb, J., Hirsch, P. E., Seidel, F., Rauter, G., & Burkhardt-Holm, P. (2022). Flow, force, behaviour: Assessment of a prototype hydraulic barrier for invasive fish. Hydrobiologia, 849(4), 1001–1019. 10.1007/s10750-021-04762-z

Wilder, M. L., Farrell, J. M., & Green, H. C. (2023). Estimating EDNA shedding and decay rates for muskellunge in early stages of development. Environmental DNA, 5(2), 251–263. 10.1002/edn3.349

Wu, Y., Colborne, S. F., Charron, M. R., & Heath, D. D. (2023). Development and validation of targeted environmental DNA (EDNA) metabarcoding for early detection of 69 invasive fishes and aquatic invertebrates. Environmental DNA, 5(1), 73–84. 10.1002/edn3.359

