## Supplementary material for "Monitoring of the invasive round goby in an estuarine seascape based on eDNA"

METHODS:

Expanded section for **3.1 dPCR assay development**

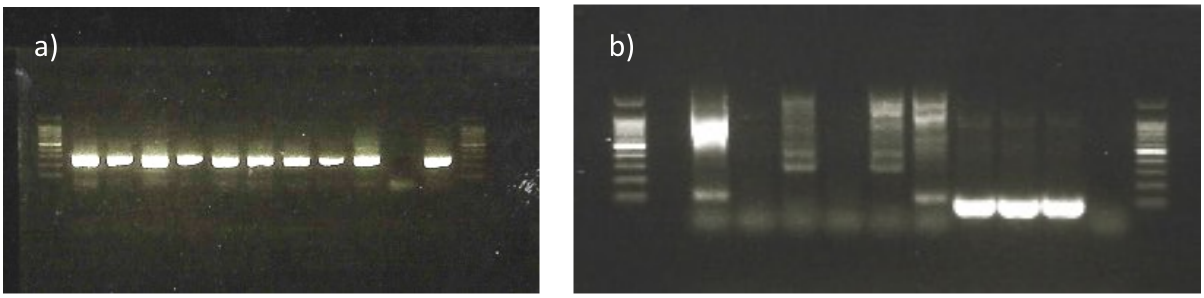


**Supplementary figure S1.** PCR results of 12S amplification with universal primers (a) and 12S assay primers (b). Sample order a: Gn-1, Nm-1, Nm-2, Pm-1, Pp-1, Pm-2, Pp-2, Gn-2, mock, water, Nm-3. Sample order b: Gn-1, Pm-1, Pp-1, Pm-2, Pp-2, Gn-2, Nm-1, Nm-2, Nm-2, mock, water.

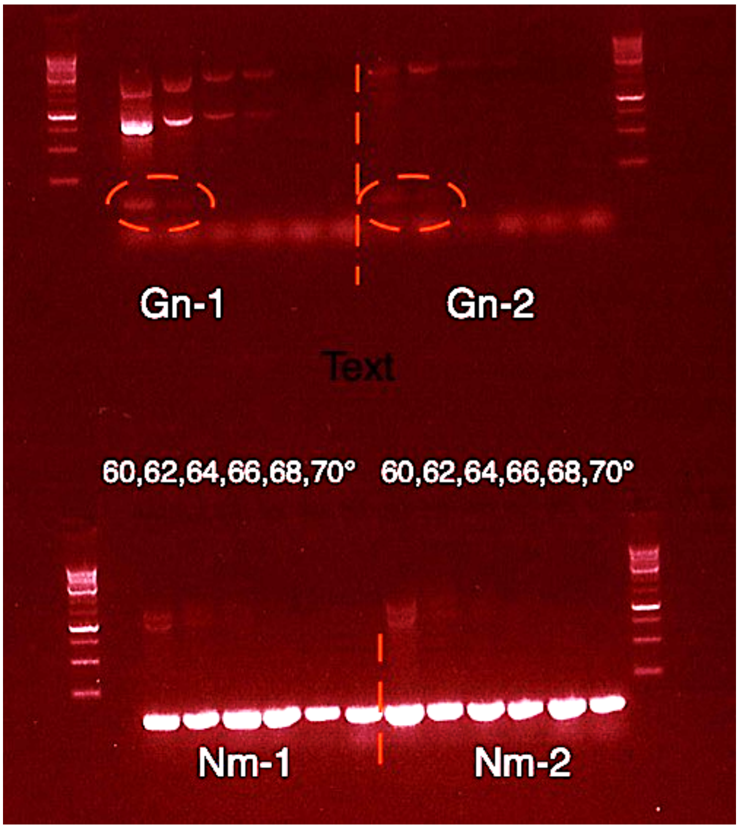


**Supplementary figure S2**. Gradient PCR with six annealing temperatures for 12S assay on the black goby (Gn) and the round goby (Nm) DNA samples. The weak bands on Gn are marked with read circles.

Expanded section for **3.4 eDNA sampling**

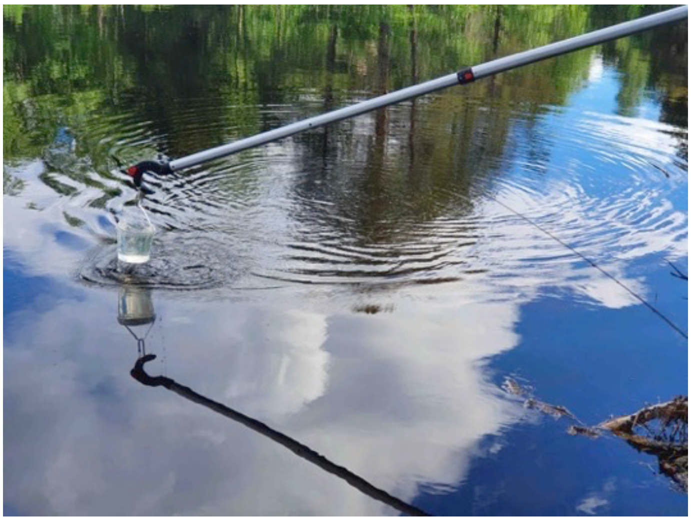

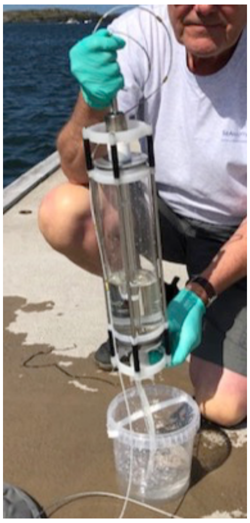

Image detailing water samplers. Boathook with clamped 1 litre sterile bucket (left) used in sampling from shallow waters (picture from another sampling) and Ruttner water sampler (right). Photo: Per Sundberg© (left), Lise-Lotte Sundberg© (right).

Expanded section for **3.6 Sampling by fishing**

Within 100 meters of each site midpoint (predetermined coordinate), three double-ended fyke nets (2 connected fykes each with a 5 meter leader, 2.63 meter funnel section with 3 funnels, with mesh ranging from 18 mm in the funnel throat to 11 mm in the cod end, REF) were placed at a depth gradient, spaced at least 15 meters apart with exact position dependent on wind direction for ease of deployment. As a complement, another 2 crab cages (plastic coated steel frame, 60 x 42 x 19 cm with entrances along both short ends and with a nylon mesh size of 13 mm) and 3 shrimp cages (circular build with diameter ~ 100 cm, 30 cm height, 10 mm nylon mesh and 3 circular openings of 8 cm diameter) were also placed at the site along the depth gradient. These were deployed in the morning to mid-day (between 08:09 and 14:00) and left until the day after (between 08:50 and 15:45) when the nets and cages were emptied. The fishing gear was deployed for an average of 26 hours and 24 minutes (Maximum 32h 26m; Mininum 22h 24 min) From the total catch, all individuals from all fish species as well as green shore crabs (*Carcinus maenas*) were counted, and all *N. melanostomus* were measured for length, and euthanized by a blow to the head followed by destruction of the brain with a sharp knife. In total 50 individuals of *N. melanostomus* were caught at 6 of the 10 sampled sites. The majority (N = 47) were caught using fyke nets, while some were also caught in the cages at 2 localities (N = 3).
